# Attractor dynamics in networks with learning rules inferred from *in vivo* data

**DOI:** 10.1101/199521

**Authors:** Ulises Pereira, Nicolas Brunel

**Affiliations:** Department of Statistics, The University of Chicago, Chicago, Illinois, USA; Department of Neurobiology, The University of Chicago, Chicago, Illinois, USA; Department of Neurobiology, Duke University, Durham, North Carolina, USA; Department of Physics, Duke University, Durham, North Carolina, USA

## Abstract

The attractor neural network scenario is a popular scenario for memory storage in association cortex, but there is still a large gap between models based on this scenario and experimental data. We study a recurrent network model in which both learning rules and distribution of stored patterns are inferred from distributions of visual responses for novel and familiar images in inferior temporal cortex (ITC). Unlike classical attractor neural network models, our model exhibits graded activity in retrieval states, with distributions of firing rates that are close to lognormal. Inferred learning rules are close to maximizing the number of stored patterns within a family of unsupervised Hebbian learning rules, suggesting learning rules in ITC are optimized to store a large number of attractor states. Finally, we show that there exists two types of retrieval states: one in which firing rates are constant in time, another in which firing rates fluctuate chaotically.

## Introduction

Attractor neural network models are one of the leading candidate scenarios for explaining how learning and memory occur in the cerebral cortex (Hopfield, 1982; Amit, 1992, 1995; Brunel, 2005). In these models, synaptic connectivity in a recurrent neural network is assumed to be set up in such a way that the network dynamics possesses multiple attractor states, each of which represent a particular item that is stored in memory. Each attractor state is a specific pattern of activity of the network, that is correlated with the state of the network when the particular item is presented through external inputs. The attractor property means that the network converges to the stored pattern, even if the external inputs are correlated to, but not identical, to the pattern, a necessary requirement for an associative memory model. In many of these models, the appropriate synaptic connectivity is assumed to be generated thanks to a ‘Hebbian’ learning process, according to which synaptic efficacies are modified by the activity of pre and post-synaptic neurons (Hebb, 2005).

These models have been successful in reproducing qualitatively several landmark observations in experiments on awake monkeys performing delayed response tasks (Fuster et al., 1971; Miyashita, 1988; Funahashi et al., 1989; Goldman-Rakic, 1995). In these experiments, animals are trained to perform a task in which they have to remember for short times the identity or the location of a visual stimulus. These tasks share in common a presentation period during which the monkey is subjected to an external stimulus, and a delay period during which the monkey has to maintain in working memory the identity of the stimulus, which is needed to solve the task after the end of the delay period. One of the major findings of these experiments is the observation of selective persistent activity during the delay period in a subset of recorded neurons in many cortical areas, in particular in prefrontal cortex (Fuster et al., 1971; Funahashi et al., 1989; Romo et al., 1999), parietal cortex (Koch and Fuster, 1989), inferior temporal cortex (Fuster and Jervey, 1981; Miyashita, 1988; Nakamura and Kubota, 1995) and other areas of the temporal lobe (Nakamura and Kubota, 1995). In those neurons, the firing rate does not decay to baseline during the delay period, but it is rather maintained at higher than baseline levels. Furthermore, this increase in firing rate is selective, i.e. it occurs only for a subset of stimuli used in the experiment. Selective persistent activity is consistent with attractor dynamics in a recurrent neural network, whose synaptic connectivity is shaped by experience dependent synaptic plasticity (Amit, 1995; Wang, 2001; Brunel, 2005).

The attractor network scenario was originally instantiated in highly simplified fully connected networks of binary neurons (Amari, 1972; Hopfield, 1982). While theorists have since strived to incorporate more neurophysiological realism into associative memory models, using e.g. asymmetric and sparse connectivity (Derrida et al., 1987), sparse coding of memories (Tsodyks and Feigel’Man, 1988; Tsodyks, 1988), online learning (Mézard et al., 1986; Parisi, 1986; Amit and Fusi, 1994), spiking neurons (Amit and Brunel, 1997; Brunel and Wang, 2001; Lansner, 2009), there is still a large gap between these models and experimental data. First, none of the existing models use patterns whose statistics is consistent with data. Most models use bimodal distributions of firing rates, with neurons either ‘activated’ by a stimulus or not, while there is no indication of such a bimodality in the data. Second, the connectivity matrix used in these models are essentially engineered (and sometimes highly fine-tuned) such as to produce attractor dynamics, but are totally unconstrained by data. Third, the attractor network scenario has been challenged by the observation of a high degree of irregularity and strong temporal variations in the firing rates of many neurons, which seem hard to reconcile with fixed point attractors (Druckmann and Chklovskii, 2012; Barak et al., 2013; Murray et al., 2017).

A recent study (Lim et al., 2015) provides us with the tools to potentially bridge these gaps. It used data from experiments in which neuronal activity is recorded in IT cortex in response to large sets of novel and familiar stimuli (Woloszyn and Sheinberg, 2012). The distribution of neuronal responses to novel stimuli permits to infer the distribution of firing rates of neurons in stimuli that are being memorized. This distribution is close to a lognormal, at odds with bimodal distributions of firing rates used in the vast majority of theoretical studies (for a few exceptions, see Treves (1990a,b); Festa et al. (2014)). Comparison between the distributions of responses to novel and familiar stimuli allows to infer the dependence of the learning rule on post-synaptic firing rates. The inferred learning rule is Hebbian, but shows two major differences with classic rules such as the covariance rule (Sejnowski, 1977):(1) The post-synaptic dependence of the rule is dominated by depression, such that the vast majority of external inputs leads to a net decrease in total synaptic inputs to a neuron with learning, leading to a sparser representation of external stimuli; (2) The dependence of the rule on post-synaptic firing rates is highly non-linear.

These results beg the question of whether associative memory can emerge in networks whose distributions of firing rates and learning rules are consistent with data. We therefore set out to study a recurrent network model in which distributions of external inputs, single neuron transfer function and learning rule are all inferred from ITC data (Lim et al., 2015). We show that: (1) learning rules inferred from visual responses in ITC lead to attractor dynamics, without any need for parameter adjustment or fine tuning; (2) Activity in the delay period is graded, with broad distributions of firing rates; (3) Learning rules inferred from data are close to maximizing the number of stored patterns, in a space of unsupervised Hebbian learning rules with sigmoidal dependence on pre and post-synaptic firing rates; (4) In a large parameter region, our model presents irregular temporal dynamics during retrieval states that strongly resembles the temporal variability observed during delay periods. In this region, retrieval states are chaotic attractors that maintain a positive overlap with the corresponding stored memory, and the network performs as a associative memory device with fluctuations internally generated by the chaotic dynamics.

## Results

We model local cortical circuits in IT cortex by a recurrent network composed of ‘firing rate’ units (Wilson and Cowan, 1972; Hopfield, 1984). The network is composed of *N* neurons whose firing rates are described by analog variables *r*_*i*_, where *i* = 1, 2, *…, N* represents the neuron index, as a simplified model for a local network in ITC (see Fig. 1 for a schematic depiction of the network). Firing rates obey standard Wilson-Cowan type equations (Wilson and Cowan, 1972)

**Figure 1:**
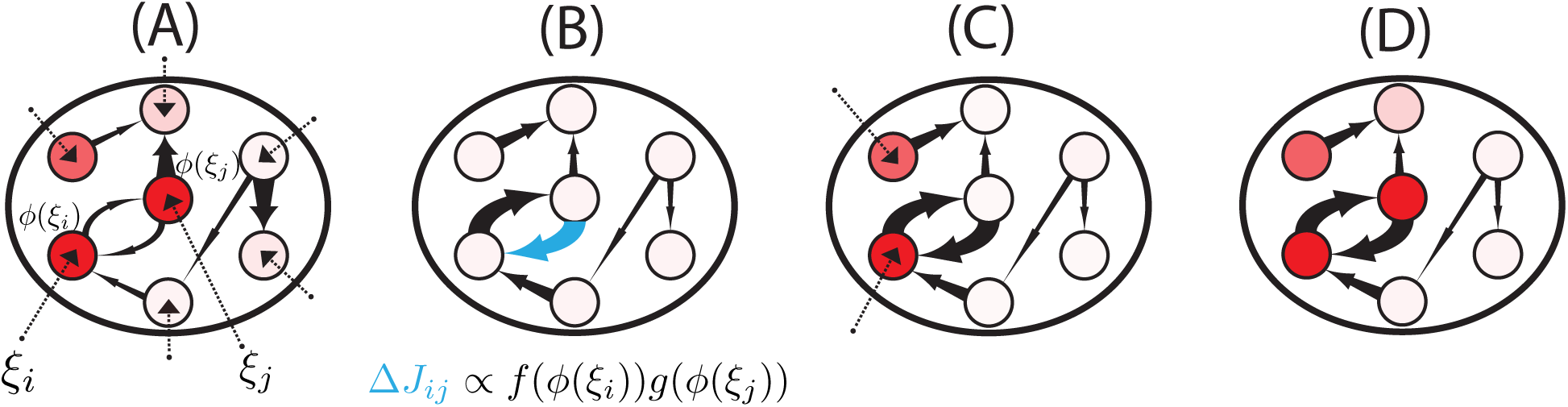
Learning and retrieval in recurrent neural networks with unsupervised Hebbian learning rules. (**A**) When a novel pattern is presented to the network, synaptic inputs to each neuron in the network (*ξ*_*l*_, for neurons *l* = 1, *…, N*) are drawn randomly and independently from a Gaussian distribution. Synaptic inputs elicit firing rates through the static transfer function, i.e. *ϕ*(*ξ*_*l*_). Some neurons respond strongly (red circles), others weakly (white circles). (**B**) The firing rate pattern produced by the synaptic input currents modifies the network connectivity according to an unsupervised Hebbian learning rule. The connection strength is represented by the thickness of the corresponding arrow (the thicker the arrow the stronger the connection). (**C**) After learning, a pattern of synaptic inputs that is correlated but not identical to the stored pattern is presented to the network. (**D**) Following the presentation, the network goes to an attractor state which strongly overlaps with the stored pattern (compare with panel A), which indicates the retrieval of the corresponding memory.

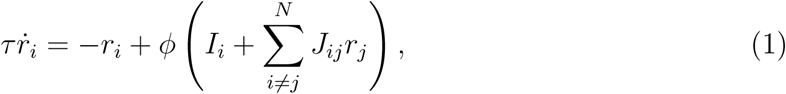

where *τ* is the time constant of firing rate dynamics, *ϕ* is the input-output single neuron transfer function (or f-I curve), *I*_*i*_ are the external inputs to neuron *i*, and *J*_*ij*_ is the strength of the synapse connecting neuron *j* to neuron *i*.

The connectivity matrix is sparse, and existing connections are shaped by external inputs (‘patterns’) through a non-linear unsupervised Hebbian synaptic plasticity rule,

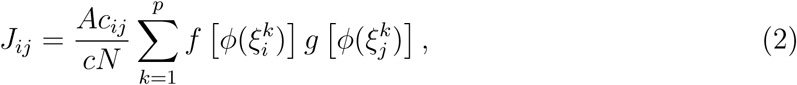

where *c_ij_* is a sparse random (Erdos-Renyi) structural connectivity matrix (c_ij_ = 1 with probability *c, c_ij_* = 0 with probability 1 c, where *c* ≪ 1), 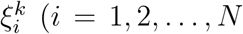 and *k* = 1, 2, …, *p*) are the external synaptic inputs to neuron *i* during presentation of pattern k, generated randomly and independently from a Gaussian distribution (see Fig. 1A,B and Methods). The external inputs shape the connectivity matrix through the firing rates 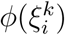 generated by such inputs, and through two non-linear functions *f* and *g* that characterize the dependence of the learning rule on the post-synaptic rate (*f*) and pre-synaptic rate (*g*), respectively. Note that this rule is a generalization of Hebbian rules used in classic models such as the Hopfield model (Hopfield, 1982) or the Tsodyks-Feigel’mann model (Tsodyks and Feigel’Man, 1988), with two important differences: patterns have a Gaussian distribution instead of binary; and the dependence of the rule on firing rates is non-linear instead of linear. In the following, the patterns that have shaped the connectivity matrix will be termed ‘familiar’ while all other random patterns presented to the network will be termed ‘novel’.

### Inferring transfer function and learning rule from data

The model defined by Eqs. (1,2) depends on three functions *ϕ*, *f* and *g* that define the single neuron transfer function and synaptic learning rule, respectively. How to choose these functions? We used a method that was recently introduced by Lim et al. (2015) to infer the tranfer function (*ϕ*) and the post-synaptic dependence of the learning rule *f* from electrophysiological data recorded in ITC (Woloszyn and Sheinberg, 2012). The transfer function *ϕ* is obtained by finding the function that maps a standard Gaussian distribution to the empirical distribution of visual responses of neurons to a large set of novel stimuli (see Methods). The post-synaptic dependence of the learning rule *f* was obtained from the differences between the distribution of visual responses to familiar and novel stimuli, under the assumption that changes in such distributions are due to changes in synaptic connectivity in recurrent ITC circuits. As an additional step to the procedure described by Lim et al. (2015), we fitted the resulting functions *ϕ* and *f* using sigmoidal functions (see Methods and Fig. 2). These sigmoidal functions provided good fits to the data (see Fig.2A-C, that shows fits of three representative ITC neurons; and Fig. S1-3 for all neurons in the data set). This fitting procedure gave us for each neurons three parameters of the transfer function: the maximal firing rate whose median value was *r*_*m*_ = 76.2 Hz, the slope at the inflection point whose median value was *β*_*T*_ = 0.82, and the threshold (current at the inflection point) whose median value was *h*_0_ = 2.46 (see Fig. 2D for a boxplot of these parameters). It also gives us for each neuron three parameters characterizing the function *f*:the threshold *x*_*f*_ (median:26.6 Hz), slope *β*_*f*_ (median: 0.28 s) and saturation *q*_*f*_ (median: 0.83). Finally, the fitting procedure also gives us the learning rate *A* (median: 3.55).

**Figure 2:**
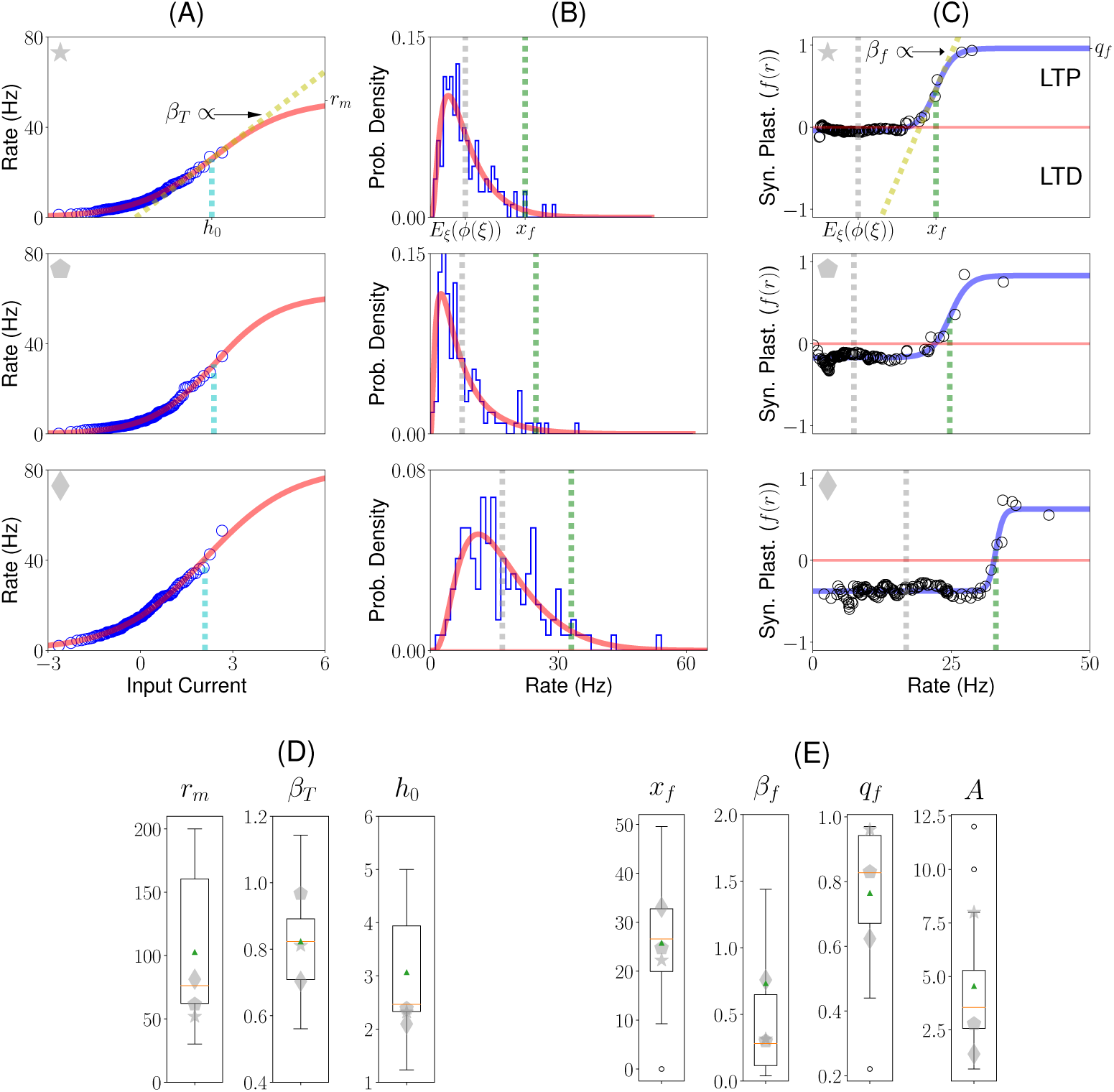
Inferring transfer function and learning rule from ITC data. (**A**) Static transfer function *ϕ* derived from the distribution of visual responses for novel stimuli, assuming a Gaussian distribution of inputs (see (Lim et al., 2015) and Methods) for three different ITC neurons. The data (blue circles) was fitted using a sigmoidal function (red line; see Methods, Equation 3), defined by three parameters: the current h0 that leads to half the maximal firing rate (cyan dashed lines), a slope parameter *β*_*T*_ (dashed yellow line in top plot), and maximal firing rate *r*_*m*_. (**B**) Distributions of firing rates in response to novel stimuli, for the same three neurons shown in A. Blue histogram: histogram of experimentally recorded visual responses. Red: Distribution of firing rates obtained from passing a standard normal distribution through the sigmoidal transfer function shown in A. Gray vertical line: average firing rate. Green vertical line: learning rule threshold *x*_*f*_ (see C) (**C**) Dependence of the synaptic plasticity rule on the postsynaptic firing rate as a function of firing rate (i.e. *f* (*r*)). The data (black circles) was fitted with a sigmoidal function (blue line; see Methods, Eq. 8), defined by three parameters: maximum potentiation *q*_*f*_; threshold *x*_*f*_ (see green dashed line); and slope parameter *β*_*f*_ (dashed yellow line in top plot). On the right axis is indicated the maximum potentiation of the fit *q*_*f*_. (**D**) Boxplot for the fitted parameters rm, *β*_*T*_ and *h*_0_ of the transfer function. (**E**) Boxplot for the fitted parameters *x*_*f*_, *β*_*f*_, *q*_*f*_ of the dependence of the synaptic plasticity rule on the postsynaptic firing rate, and A, the learning rate. The red line and green triangle indicate the median and the mean of the fitted parameters, respectively. Gray symbols indicate the parameters of the three neurons shown in A,B,C.

A number of features of these fitted functions are noteworthy: First, the vast majority of the visual responses of neurons are in the supralinear part of the transfer function, and therefore far from saturation. This is consistent with many studies showing supra-linear transfer functions at low firing rates, both *in vitro* (Rauch et al., 2003) and *in vivo* (Anderson et al., 2000). Second, this has the consequence that the distribution of visual responses are strongly right-skewed, and in fact close to lognormal distributions, consistent with multiple observations *in vivo* (Hromadka et al., 2008; Roxin et al., 2011; Buzsaki and Mizuseki, 2014; Lim et al., 2015). Third, the function *f* presents two major differences compared to standard Hebbian rules such as the covariance rule (Sejnowski, 1977; Tsodyks and Feigel’Man, 1988): It is strongly non-linear; and the threshold between depression and potentiation occurs at a firing rate that is much higher than the mean rate, leading to depression of the mean synaptic inputs to a neuron for the vast majority of shown stimuli. Consequently, the average of the function *f* across the distribution of patterns is negative, which leads to a decrease of the average visual response with familiarity (Lim et al., 2015).

The only parameters that are left unconstrained by data are two parameters characterizing the function *g*. In most of the following, we will take those parameters to be identical to the corresponding parameters of the function *f* (i.e. *x*_*g*_ = *x*_*f*_ and *β*_*g*_ = *β*_*f*_; note that *q*_*g*_ is fixed by the condition that the average of the function *g* across the distribution of patterns is zero, see Methods). We will also explore the space of values of *x*_*g*_ and *β*_*g*_ (see below).

### Dynamics of the network following presentation of a familiar stimulus

Having specified the model, we now turn to the dynamics of the network described by Eqs. (1,2), whose parameters are set according to the procedure described above. In particular, we ask whether the model exhibits attractor dynamics. To address this question, we used both numerical simulations of large networks (see methods) and mean field theory (MFT see methods and mathematical note on SI). For the MFT, we assume that both the number of neurons and stored patterns are large (i.e. more specifically the limit *p, N→∞*), while the number of stored patterns *p* divided by the average number of synapses per neuron (*Nc*),α ≡ *p/Nc* remains of order one. We call *α* the *memory load* of the network. The results of the MFT only depend on *N*, *c* and *p* via this quantity (see methods and mathematical note on SI). From our MFT analysis, we obtain mathematical expressions for two ‘order parameters’ that describe how network states are correlated (or not) with stored patterns. We are specifically interested here in the situation when the network state is correlated with one of the stored patterns (e.g. following the presentation of this particular pattern).

The first order parameter describes the ‘overlap’ *m* between the current state of the network (described by the vector of firing rates *r*_*i*_, for *i* = 1, 2, *…, N*) and the pattern of interest (see Methods for the mathematical definition of *m*). When *m* is of order 1, this indicates that the corresponding pattern is retrieved from memory. Consequently, each pattern stored in memory can be retrieved by initializing the network dynamics with a configuration that is close to that particular pattern, and letting the network evolve towards its attractor state. In this case, giving a partial cue to the network leads to the dynamics towards an attractor state correlated with the stored pattern, a signature of associative memory. The other order parameter *M* describes the interference due to the other stored patterns in the connectivity matrix; it is proportional to the average squared firing rates of the network (see Methods). Equations for the order parameters as a function of, α, *ϕ*, *f* and *g* are given in Methods.

The results of the simulation of a particular realization of a network of *N* = 50, 000 neurons with *c* = 0.005 (an average of 250 connections per neuron) storing *p* = 30 patterns (α = 0.12), and the comparison with the results from MFT are shown in Fig. 3. In the simulations, the network was initialized in a state which was uncorrelated with all the stored patterns. For these parameters, the network converged to a ‘background’ state in which all neurons fire at low rates (average 7.98/s, standard deviation 2.92/s). Upon presentation of a novel stimulus (Fig. 3A), neurons were driven to stimulus-specific firing rates, with a distibution of firing rates that was close to a lognormal distribution (Fig. 3C), similar to experimental observations (Lim et al., 2015). The distribution is close to lognormal because the distribution of inputs to neurons is Gaussian, and the neuronal transfer function is close to being exponential at low rates (see Methods). After the end of the presentation of the stimulus, the network came back to its initial background state (Fig. 3A). Upon presentation of a familiar stimulus (Fig. 3D), the statistics of neuronal responses differed markedly from the response to novel stimuli: a few neurons responded at higher rates, but the majority of neurons responded at lower rates compared to a novel stimulus. The distribution of visual responses for familiar stimuli had consequently a lower mean compared to the distribution of responses for novel stimuli but a larger tail at high rates (compare Fig. 3C and F). These two features were consistent with data recorded in ITC by multiple groups (Li et al., 1993; Kobatake et al., 1998; Logothetis et al., 1995; Freedman et al., 2006; Woloszyn and Sheinberg, 2012).

**Figure 3:**
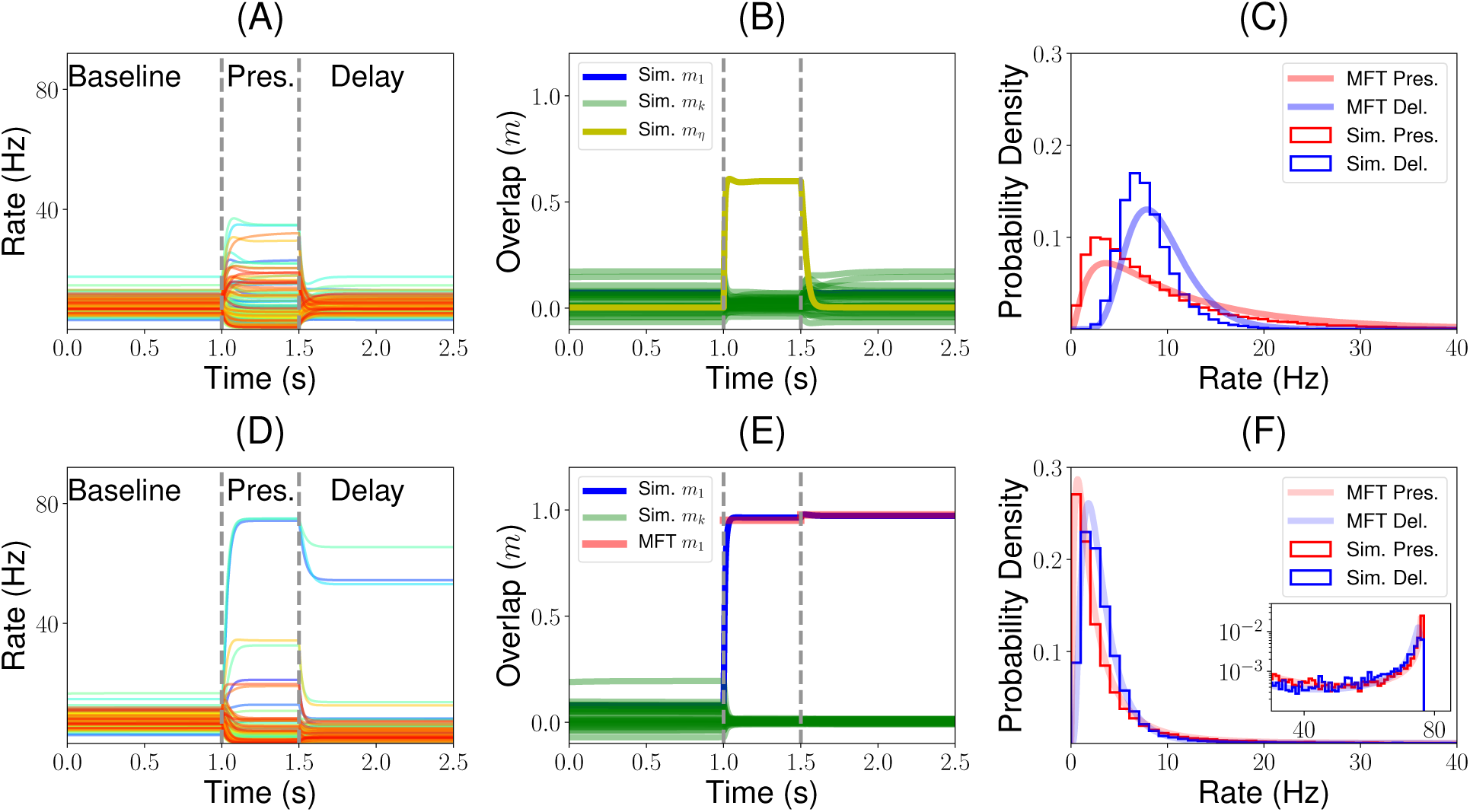
Dynamics of the network before, during and after the presentation of novel (top row) and familiar (bottom row) stimuli, mimicking the initial part of a trial of a delay match to sample (DMS) experiment. (**A**) Firing rate of a randomly sampled subset of 100 neurons of a simulated network before, during and after the presentation of a novel stimulus. Vertical dashed lines indicate the beginning and the end of the presentation. Note that the firing rates of all neurons decay to baseline following removal of the stimulus. (**B**) Dynamics of the overlaps with the stored patterns. Green traces show overlaps computed numerically from the network simulation corresponding to each of the stored patterns. The yellow trace shows the overlap of the network state with the shown novel pattern. (**C**) Distribution of firing rates during the presentation (red) and delay (blue) periods. Smooth curves correspond to the predictions of the MFT, histograms are obtained from network simulations. (**D**) Similar to A, except that the shown stimulus is familiar. Note that this time firing rates do not decay to baseline during the delay period, but to a value that is strongly correlated (but not identical) to the visual response. (**E**) Dynamics of overlaps when a familiar stimulus is presented. The blue trace shows the numerically computed overlap with the pattern presented during the presentation period. The red trace shows the corresponding overlap computed from MFT. (**F**) Distribution of firing rates during the presentation (red) and delay (blue) periods in response to the presentation of a familiar stimulus. Note that the distributions appear unimodal, with most neurons firing in the 0-10Hz range. However a zoom on the right side of the distribution shows a tiny peak close to saturation (see text).

After removal of a familiar stimulus, the network no longer came back to the initial background state, but rather converged to an attractor state that was strongly correlated with the shown stimulus (Fig. 3D), as shown by the strong overlap between the network state and the shown pattern (see blue curve in Fig. 3E). A small fraction of neurons exhibited persistent activity at high rates (4.5% of the neurons are above half maximal rate), but most neurons remained at low rates during the simulated delay period (Fig. 3F). The distribution of firing rates was again similar to a lognormal distribution at low rates, but the tail of the distribution was shaped by neuronal saturation and therefore exhibited a tiny peak close to maximal firing rates. Both overlap with presented pattern and distributions of firing rates could be computed by the MFT and were in close agreement with network simulations (Fig. 3E and F).

Thus, our network behaved as an associative memory when constrained by ITC data, without any need for parameter variation or fine tuning. Furthermore, in addition to reproducing the distributions of visual responses for both novel and familiar stimuli seen experimentally, it also exhibited qualitatively some of the main features observed both during spontaneous and delay activity in IT cortex: broad distribution of firing rates in both spontaneous and delay period activity, and small fraction of neurons firing at elevated rates during persistent activity (Miyashita, 1988; Nakamura and Kubota, 1995).

### Storage capacity, and its dependence on *g*

We now turn to the question of the storage capacity of the network, i.e. how many different patterns can be stored in the connectivity matrix. The calculation of the storage capacity of associative memory models such as the Hopfield model was one of the first successful applications of statistical physics to theoretical neuroscience (Amit et al., 1987). One of the main findings of such models is that the number of patterns that can be stored scales linearly with the number of plastic connections per neuron, i.e. the maximal value of, α is of order 1. This maximal storage capacity, α_*c*_ has been computed in many variants of the Hopfield model (see e.g. Amit (1992)). To compute the storage capacity of our network, we found numerically the largest value of, α for which retrieval states (i.e. states with positive overlap with one of the stored patterns, *m >* 0) exist. Fig. 4A shows how the overlap in retrieval states *m* varies as a function of the storage load, α, computed using both MFT (solid line) and simulations (symbols with errorbars) when parameters of the functions *ϕ* and *f* are taken to be the median best-fit parameters, and those of the function *g* are taken to be identical to *f*. It shows that *m* gradually decreases with, α, due to more ‘noise’ in the retrieval due to other stored patterns, until it drops abruptly to zero at a value of, α_*c*_ = 0.56. This value is remarkably similar to the maximal capacity of the sparsely connected Hopfield model of binary neurons storing binary patterns, for which, α_*c*_ = 0.64 (Derrida et al., 1987).

**Figure 4:**
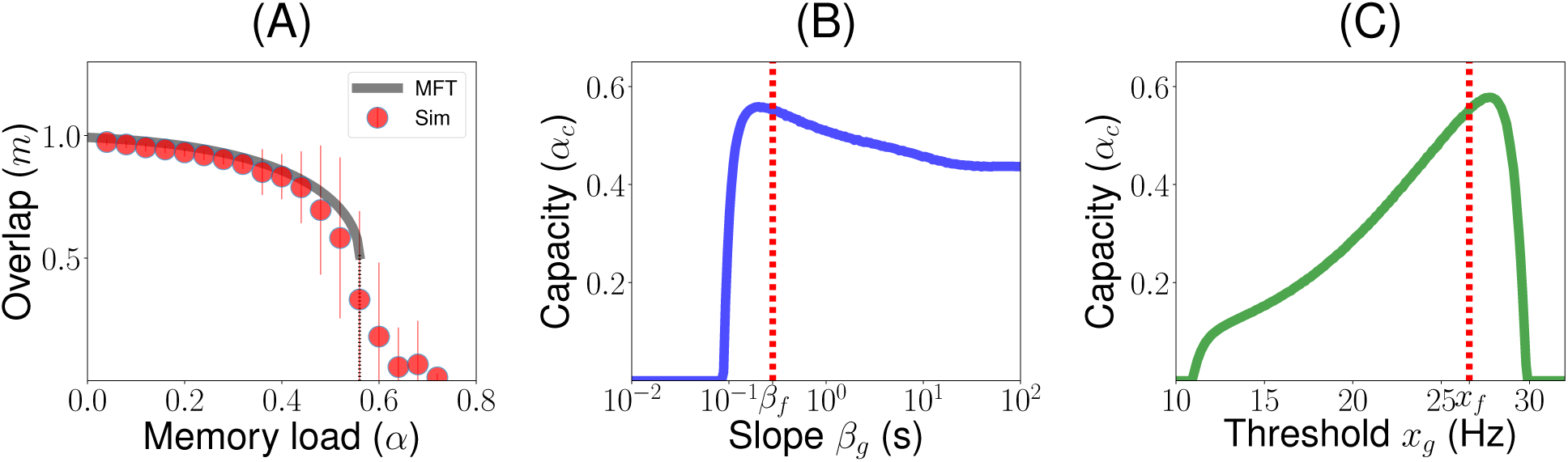
Storage capacity of the network, and its dependence on *g*. (**A**) Overlap as a function of memory load, α (number of patterns stored divided by average number of connections per neuron). Grey: MFT. Red circles: Numerical simulations (average and standard deviations computed from 100 realizations with *N* = 5 *·* 10^4^). The overlap stays positive until *α ∼* 0.56. Parameters of *g* are chosen to be identical to those of *f*. (**B**) Capacity vs *β*_*g*_. The capacity is maximized for *β*_*g*_ *∼ β*_*f*_ (dashed red line *β*_*g*_ = *β*_*f*_). (**C**) Capacity vs *x*_*g*_. The capacity is close to being maximized for *x*_*f*_ *∼ x*_*g*_ (dashed red line *x*_*g*_ = *x*_*f*_). Other parameters as in Fig. 3.

We then explored how the capacity depends on the parameters of the function *g*, that describes the dependence of the learning rule on the presynaptic firing rate. Fig. 4B and C show that the capacity is close to being maximized when these parameters match those of the function *f*, i.e. *x*_*g*_ = *x*_*f*_ and *β*_*g*_ = *β*_*f*_. Fig. 4B shows that the capacity is non-zero only when the *g* is sufficiently non-linear, i.e. *β*_*g*_ *>* 0.1. It peaks around *β*_*g*_ = *β*_*f*_, but remains high in the *β → ∞* limit when the function *g* becomes a step function. Fig. 4C shows that the capacity is non-zero only in a finite range of *x*_*f*_, between 10 and 30/s. It shows again that capacity peaks when *x*_*g*_ is close to *x*_*f*_.

### Learning rules inferred from ITC data are close to maximizing memory storage

The storage capacity of the network with median parameters is in the same range or higher than the capacity of classic associative memory models of binary neurons for instance, the Hopfield model has a capacity of *α*_*c*_ *∼* 0.14 (Amit et al., 1987), while its sparsely connected variant has a capacity of *α*_*c*_ ∼ 0.64 (Derrida et al., 1987). The next question we addressed is how this capacity depends on the parameters of this learning rule. We have already discussed above the dependence of the capacity on *x*_*g*_ and *β*_*g*_. Here, we explore the dependence on the four remaining parameters characterizing the learning rule *A*, *x*_*f*_, *β*_*f*_ and *q*_*f*_. Using MFT, we explored systematically the space of these four parameters, and plot in Fig. 5 all possible cuts of this four dimensional space, in which 2 of the 4 parameters are varied, while the other 2 are set to the median values. In all these plots, the maximal capacity *α*_*c*_ is plotted as a function of two parameters, using a gray scale (white indicate high capacity, black low capacity). The yellow dashed line indicates the line for which the function *f* is ‘balanced’ (i.e. its average across the distribution of patterns is zero). It marks the border between a depression-dominated region, for which learning leads to a decrease in average responses, and a potentiation-dominated region, for which learning leads to an increase of such responses. The red cross mark indicates the median parameters, while the dashed red rectangle indicate the interquartile range.

**Figure 5:**
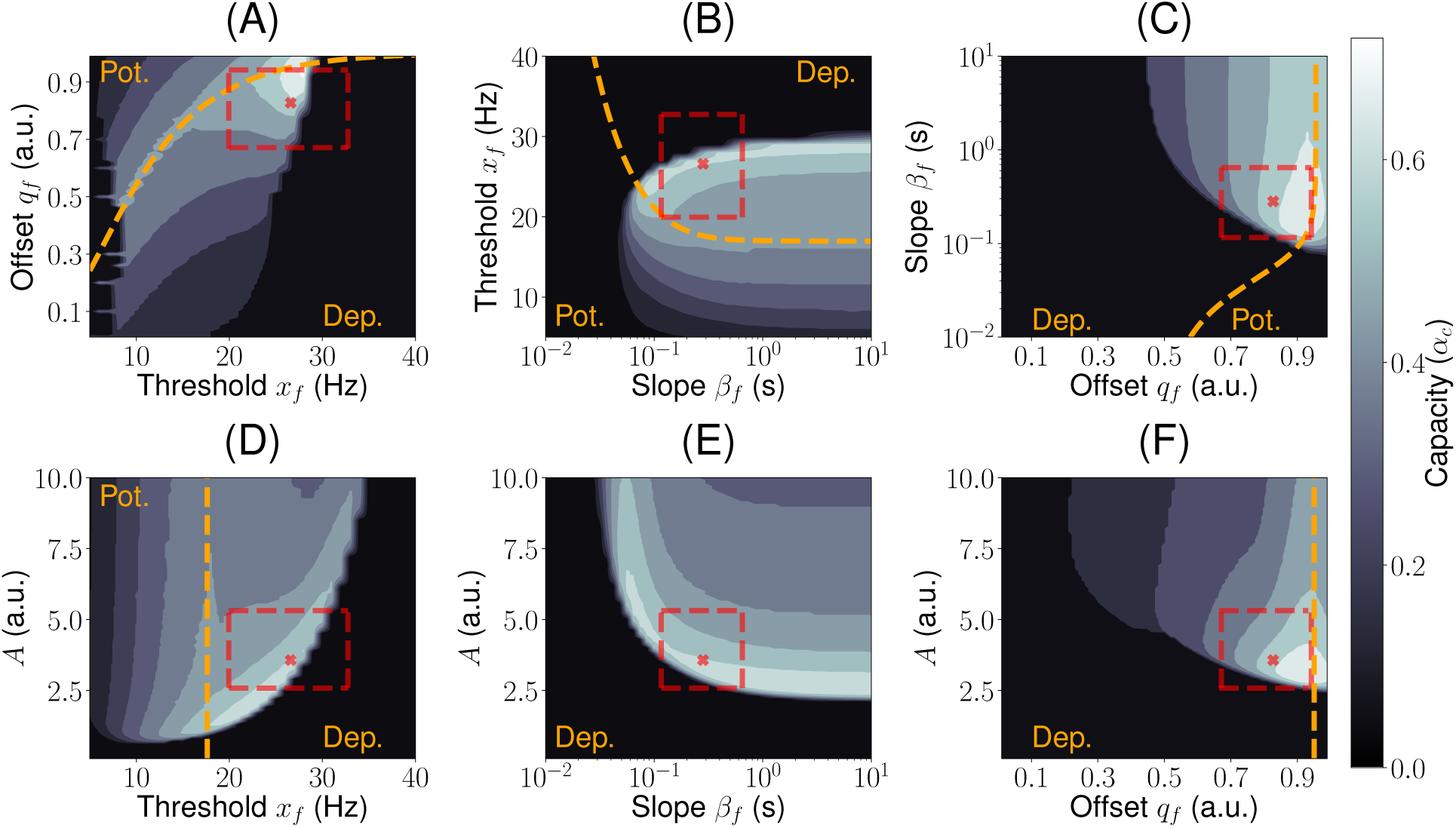
Inferred learning rules from ITC are close to maximizing memory storage. Contour plots for the capacity of the network as a function of two parameters. In each plot, two parameters are set to the median best-fit parameters, and the other two are varied. The yellow dashed line indicates the curve where potentiation and depression are balanced in average 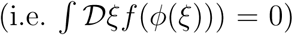. It separates the potentiation 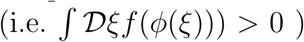 and depression 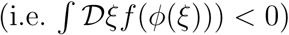 regions. The parameter region corresponding to the IQR is indicated with a red dashed rectangle. The median bestt parameters are shown as a red cross mark. The parameters of g: *x*_*g*_ = *x*_*f*_ and *β*_*g*_ = *β*_*f*_.

Fig. 5 shows that the median parameters are close to maximizing storage capacity. In fact, we found that the maximal capacity over this space is *α*_*c*_ *≈*0.85 (see Fig. S4 and SI Mathematical Note for details). These figures show also that most (but not all) of the interquartile range lie in a high-capacity region. It also shows that some parameter variations lead to little changes in capacity, while others lead to a drastic drop. Decreasing the learning strength *A* from its optimal value leads to an abrupt drop in capacity, while increasing it leads to a much gentler decrease (see Fig. 5D-F). A similar effect is observed for the slope of *f*; decreasing the slope (i.e. making *f* more linear) leads to an abrupt decrease in capacity, while increasing it beyond the median value leads to very little change in capacity (see Fig. 5B-D). Thresholds *x*_*f*_ for which high capacities are obtained are much higher than the mean visual response (Fig. 5A,B and D), leading to a sparsening of the representations of the patterns by the network. Finally, the optimal offset is close to the ‘balanced’ line, but slightly on the depression-dominated region, as the median parameter (Fig. 5A,C and F).

### A chaotic phase with associative memory properties

Are fixed point attractors the only possible dynamical regime in this network? Firing rate models with asymmetric connectivity have been shown to exhibit strongly chaotic states (Sompolinsky et al., 1988; Tirozzi and Tsodyks, 1991). Varying parameters of the learning rule, we found parameter regions in which background and/or retrieval fixed point attractor states destabilize and the network settle into strongly chaotic states. Fig. 6A shows an example of such chaotic states, obtained for the median parameters as in Fig. 3, except for the learning rate (*A* = 11.5). For such parameters, the background state is strongly chaotic. Presentation of a familiar stimulus leads to a transition to another chaotic state, in which all neurons fluctuate chaotically around stimulus-specific firing rates, such that the mean overlap with the corresponding pattern remains high (see Fig. 6 B). Remarkably, chaotic retrieval states remain strongly correlated with the corresponding patterns (see Fig. 6B), so that the network can still perform as an associative memory in spite of the chaotic fluctuations of network activity. Interestingly, the storage capacity for such parameters is larger than the capacity estimated from the static MFT (see Fig. 6C).

**Figure 6:**
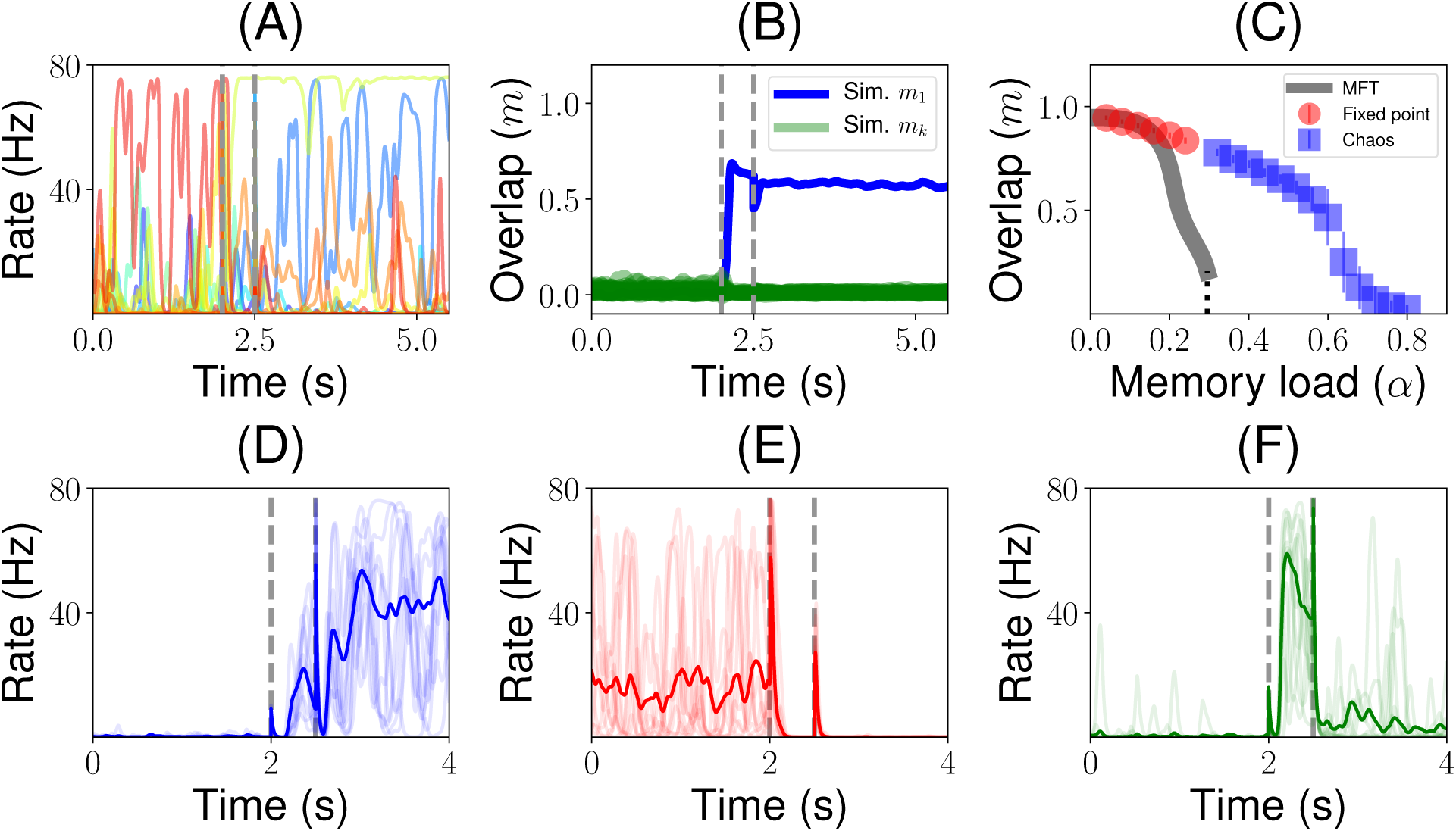
Chaotic background and retrieval states, for a network with different parameters as in Fig. 3. (**A**) Firing rate dynamics for a randomly sampled subset of 10 neurons of a simulated network when a familiar stimulus (i.e. one of the stored patterns) is presented.(**B**) Dynamics of the overlap during the presentation of a familiar stimulus. Green traces shown all the overlaps computed numerically from the network simulation corresponding to each of the stored patterns except the one with the presented pattern, shown in blue. (**C**) Overlap vs memory load. Gray curve: MFT. Red circles: simulations in which the dynamics converge to fixed point attractors. Blue square: simulations in which the dynamics converge to chaotic states. (**D**-**F**) Dynamics of the firing rate of three example neurons in 10 different trials (random initial conditions transparent traces). Trial-averaged firing rate (over 20 trials) is shown with an opaque trace. Parameters as in Fig. 3, except for the learning rate (*A* = 11.5) and memory load (*α* = 0.48 in all panels except in C).

In such chaotic retrieval states, single neuron activity exhibit strong firing rate fluctuations which vary from trial to trial (see thin colored lines in Fig. 6D-F showing three randomly selected neurons), but trial-averaged firing rates show systematic temporal patterns. For instance, the activity of the neuron shown in Fig. 6D ramps up in the first second of the delay period, before this activity plateaus at a rate of about 40/s. The neuron shown in Fig. 6F shows a rapid activity increase during the presentation period, but then this activity drops to an intermediate rate of about 5/s during the delay period. These temporal patterns of the trial-averaged firing rate, together with a strong irregularity within trials, are remininscent of observations by multiple groups in primate PFC during delay periods (Shafi et al., 2007; Brody et al., 2003; Murray et al., 2017).

## Discussion

We have shown that a learning rule inferred from data generate attractor dynamics, without any need for parameter adjustment or tuning. Furthermore, this rule produces a storage capacity that is close to the maximal capacity, in the space of unsupervised Hebbian learning rules with sigmoidal dependence on both pre and post-synaptic firing rates. Remarkably, as the inferred learning rules from ITC recordings, learning rules that maximize memory storage depress the bulk of the distribution of the learned inputs (those that lead to low to intermediate firing rates) while potentiating outliers (those that lead to high rates), leading to a sparse representation of stored memories. The attractor states generated by our model are characterized by graded activity with a continuous range of firing rates (Treves, 1990a,b; Festa et al., 2014). Most of the distribution lies in the low rate region of the neuronal transfer function, leading to a strongly skewed distribution, with a small fraction of neurons firing at higher rates. These observations are consistent with the available data in ITC during delay match to sample experiments (Miyashita, 1988; Nakamura and Kubota, 1995).

For a range of parameters values consistent with learning rules inferred from data, our model presents irregular temporal dynamics for retrieval states, similar to the temporal and across trial variability observed during delay periods in multiple studies (Murray et al., 2017). In this regime, retrieval states are chaotic, yet they maintain non-zero overlap with the corresponding memories. Thus, the network performs robustly as an associative memory device, even though strong fluctuations are internally generated by its own chaotic dynamics.

### Distribution of firing rates

Our model naturally gives rise to highly skewed distributions of firing rates, consistent with those that have been observed during presentation of visual stimuli in ITC (Lehky et al., 2011; Lim et al., 2015) and during delay periods of DMS tasks (Miyashita, 1988; Nakamura and Kubota, 1995). It also reproduces the decrease in the mean response with familiarity, and the increase in selectivity with familiarity. Our model shows for most of the explored parameter space a weak bimodality in the distribution of firing rates due to neuronal saturation in response to familiar stimuli, with a tiny peak close to neuronal saturation. Detecting this bimodality in data would require far larger stimuli sets than have been used so far, which might be difficult given experimental constraints. We also note that the shape of the transfer function close to saturation is only weakly constrained by data, since most of the data points lie in the supralinear range of this transfer function. Transfer functions with different behaviors at high rates will have an effect on the tails of the distributions. We have checked for instance that a transfer function with a square root behavior at high rates (Brunel, 2003) leads to a broader peak at high rates.

### Learning rule

The learning rule we have used in our network model was inferred from ITC data (Lim et al., 2015). It is an unsupervised Hebbian rule, as it only depends on the pre and postsynaptic firing rates, and it leads to potentiation for large pre and post-synaptic rates. As other popular examples of Hebbian rules such as the covariance rule (Sejnowski, 1977)or the the BCM rule (Bienenstock et al., 1982), it is separable in pre and post-synaptic rates. It also reproduces some of the phenomenology of the dependence of synaptic plasticity on pre and post-synaptic firing rates in cortical slices; in particular, large pre and postsynaptic firing rates lead to LTP (Sjöström et al., 2001). Large pre-synaptic firing rate in conjunction with low post-synaptic firing rate, lead to depression, consistent with ‘pairing’ experiments in which LTD is triggered by pre-synaptic activity, together with intermediate values of the membrane potential (Ngezahayo et al., 2000). Plasticity at low pre-synaptic firing rates could be due to plasticity mechanisms leading to ‘normalization’ or homeostasis.Indeed, our plasticity rule could be written as 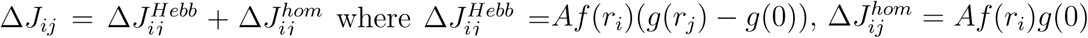. The homeostatic’ component 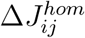 leads to a decrease in the efficacy of all synapses onto a post-synaptic neuron when the neuron is firing at high rates, while it leads to an increase when the neuron fires at low rates (note g(0) < 0).

To derive the learning rule, we used a subset of the data recorded by Woloszyn and Sheinberg (2012), i.e. excitatory neurons that show negative changes at low rates and positive changes at high rates. Those neurons are approximately half (14/30) of the neurons that showed significant differences between the distributions of visual responses for familiar and novel stimuli. Out the remaining 16 neurons, 10 showed negative changes for all rates, while 6 showed the opposite pattern of positive changes for all rates. This heterogeneity in inferred learning rules could be due to a heterogeneity in neuronal properties for instance, it could be that the ‘putative’ excitatory neurons recorded in this study form a heterogeneous group of cells, some of which might actually be inhibitory.Consistent with this, some inhibitory neuron classes have electrophysiological properties (and in particular, spike width) that are closer to pyramidal cells that to fast-spiking interneurons. Another possibility is that part of the apparent heterogeneity stems form the same underlying learning rule, but with heterogeneous parameters. For instance, inferred learning rules with negative changes at all rates are consistent with a sigmoidal post-synaptic dependence *f*, but with a high threshold *x*_*f*_ that lies above the range of firing rates elicited in that particular experiment. Elucidating which of these scenarios hold in IT cortex will need recordings from more neurons, as well as recordings of single neurons with more stimuli.

Our approach is complementary to other studies that have inferred learning rules from *in vitro* studies, and then shown that these rules lead to attractor dynamics in large networks of spiking neurons (Litwin-Kumar and Doiron, 2014; Zenke et al., 2015). In contrast to these studies, we showed that a network with learning rules inferred from *in vivo* data can achieve a high storage capacity, and generate graded distributions of firing rates during visual presentation and delay periods. It will be interesting to investigate whether, and in which conditions spike-timing and voltage based learning rules used in such studies can produce a firing rate dependence that is consistent with the rule used here.

### Time-varying neural representations

In recent years, the standard attractor network scenario has been challenged by multiple observations of strong variability and non-stationarity during the delay period in prefrontal cortex (Compte et al., 2003; Shafi et al., 2007; Barak et al., 2010; Barak and Tsodyks, 2014; Kobak et al., 2016; Murray et al., 2017). Statistical analysis of recordings in this area during two different working memory tasks has shown that variability observed during delay periods is consistent with static coding of the stimulus kept in memory (Murray et al., 2017). Various models have been proposed to account for variability and/or non-stationarity (Barbieri and Brunel, 2007; Mongillo et al., 2008; Lundqvist et al., 2010; Mongillo et al., 2012; Druckmann and Chklovskii, 2012).

Here we propose an alternative mechanism where chaotic attractors with associative memory properties naturally generate the time-varying irregular activity observed during delay periods in associative memory tasks. In this state, chaotic attractors correspond to internal representations of stored memories. Each chaotic attractor state maintains a positive overlap with the corresponding stored memory. In this scenario, the network performs as an associative memory device where temporal variability is generated internally by chaos. This model naturally exhibits the combination of strong temporaly dynamics yet stable memory encoding which has been demonstrated in PFC by various groups (Druckmann and Chklovskii, 2012; Murray et al., 2017). It will be interesting to compare this model to existing data, using for instance methods used in Murray et al. (2017).

There has been a longstanding debate whether the type of chaotic states seen in firing rate models can be seen also in spiking network models under the form of ‘rate chaos’. Recent studies indicate that this type of chaos can be observed provided coupling is sufficiently strong, as in firing rate models Ostojic (2014); Harish and Hansel (2015); Kadmon and Sompolinsky (2015). Thus, it is reasonable to expect that the type of retrieval chaotic states we observed in our network can also be realized in networks of spiking neurons.

### Optimality criteria for information storage

Here, we have argued that learning rules that are inferred from electrophysiological recordings in ITC of behaving primates are close to optimizing information storage, in the space of unsupervised Hebbian learning rules that have a sigmoidal dependence on both pre and post-synaptic firing rates. Such learning rules are appealing because synapses do not need to know anything beyond the firing rates of pre and post-synaptic neurons to form memories, two quantities that are easily available at a synapse. However, one cannot exclude that the dependence of plasticity on neuronal activity takes other forms than the one investigated here. In particular, a potentially more powerful approach proposed by Gardner (1987) relies in maximizing the number of attractors in the space of all possible synaptic matrices. Unsurprisingly, this approach leads in general to a larger capacity than the ones that can be achieved by unsupervised Hebbian rules, but it turns out that in sparse coding limit, the covariance rule reaches asymptotically the Gardner bound (Tsodyks and Feigel’Man, 1988; Tsodyks, 1988). These results have been obtained in networks of binary neurons, and it remains to be investigated whether similar results could be obtained in networks of analog firing rate neurons. An additional challenge in comparing the two approaches in such networks is that the stored attractors are in our case not identical to the pattern that was initially shown to the network, while in the standard Gardner approach, the two were constrained to be identical.

Altogether, our results strongly reinforce the link between attractor network theory and electrophysiological data during delayed response tasks in primates. Furthermore, they suggest that learning rules in association cortex are close to maximizing the number of possible internal representations of memories as attractor states.

## Methods

### Data analysis

We reanalyze the data recorded by Luke Woloszyn and David Sheinberg (Woloszyn and Sheinberg, 2012) using the method described in Lim et al. (2015). This data consists in trial-averaged firing rates of individual neurons in ITC (in a time window between 75 ms and 200 ms after stimulus onset) in response to 125 novel and 125 familiar stimuli measured, during a passive fixation task. We focused on the 30 putative excitatory neurons whose distributions of visual responses for novel and familiar stimuli were significantly different, using the Mann-Whitney U test at 5 significance level. In these neurons, the postsynaptic dependence of the learning rule, was inferred using the method described in Lim et al. (2015). In this subset of neurons, we focused on 14 excitatory neurons, the ones that show negative input changes for low firing rates and positive input changes for high firing rates. For these 14 neurons, the transfer function *ϕ*, and the postsynaptic dependence of the learning rule, *f*, are inferred using the method described in Lim et al. (2015).

The first step is to infer the transfer function *ϕ*. We assume that inputs to neurons during presentation of novel stimuli have a Gaussian distribution. The transfer function is then obtained as the function *ϕ* that maps a standard Gaussian to the empirical distribution of firing rates for novel stimuli (Lim et al., 2015). In practice, the function is obtained by building a quantile-quantile plot between the distribution of firing rates for novel stimuli and the assumed standard normal distribution of inputs (see Fig. 2 A and B and S1-2). The obtained transfer function (blue circles in Fig. 2) was fitted with the sigmoidal function

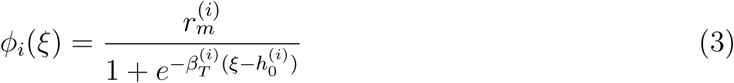

where 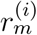 is the maximal firing rate, 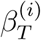 measures the slope at the inflection point, and 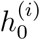 is the location of this inflection point. *h*_0_ is also the current leading to half maximal firing rate. These parameters were obtained by minimizing the squared error. We thus obtained for each of the 14 neurons the best estimators 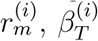 and 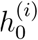 with *i* = 1, 2, …, 14 whose statistics are summarized in Fig. 2D.

The next step is to infer the postsynaptic dependence of the learning rule, *f*. For this, we use the difference between the distributions of visual responses to novel and familiar stimuli (Lim et al., 2015). In the model, learning of a novel stimulus defined by inputs 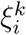 that leads to firing rates 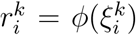 leads to changes in recurrent inputs, due to changes in synaptic inputs

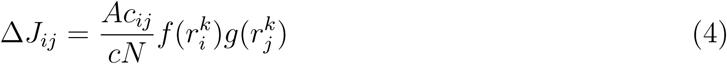

This leads to a change in total inputs to neurons that is proportional to

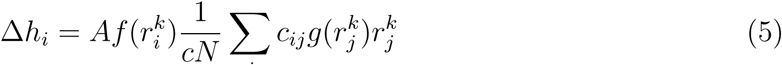

In the large *N* limit, Eq. (5) becomes

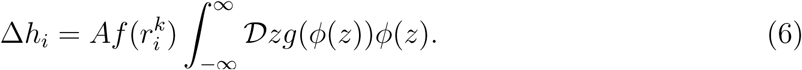

where 𝒟*z* is the standard Gaussian measure, 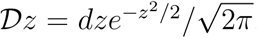. Eq. (6) give us the relationship between changes of total inputs to a neuron with learning of a particular stimulus, and the firing rate of the neuron upon presentation of that stimulus for the first time. This relationship can be inferred from the data by computing the difference between the quantile function of visual responses to familiar stimuli and the quantile function of visual responses to novel stimuli, and by plotting this difference as a function of visual response to novel stimuli (Lim et al., 2015). We then fitted the input change with a sigmoidal function given by

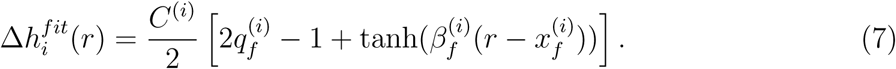

where *C*^(*i*)^ gives the amplitude of the total changes, 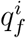 measures the vertical offset of the curve (for *q*_*f*_ = 1, Δ*h* is non-negative at all rates, while for *q*_*f*_ = 0 it is non-positive at all rates), 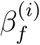 measures the slope at the inflection point, and 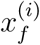 is the rate at the inflection point. In the following, we refer to 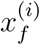 as the threshold since it is typically very close to the rate at which Δ*h* changes sign. For each of the 14 neurons, the parameters *C*^(*i*)^, 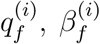 and 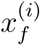 with *i* = 1, 2, …, 14 were estimated by minimizing the squared error. The inferred function *f* for each neuron is given by

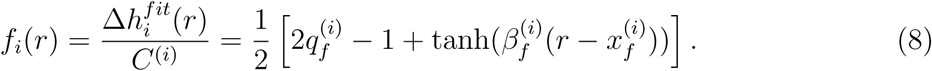

The parameter *A* is then obtained as

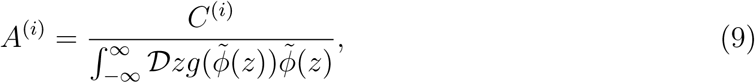

Where 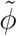 is the sigmoidal transfer function in Equation 9 whose parameters are the medians of the fitted parameters. The function *g* was also chosen to be a sigmoid, given by

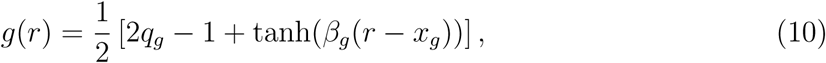

with *q*_*g*_ set such that the average change in connection strength due to learning of a single pattern is zero, i.e.

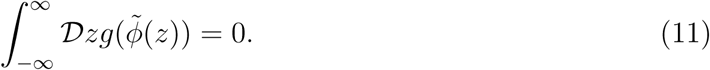

Note that *g* is unconstrained by data. For most of the paper, we set the slope and the threshold for *g* to the median of the fitted parameters for *f*, i.e. 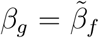 and 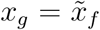. We also explored how the capacity depends on *β*_*g*_ and *x*_*g*_, as shown in Fig. 3.

### Mean field theory

Here we present the main results of our mean field analysis that quantifies the retrieval of a particular familiar pattern during the delay period. Detailed calculations for this case are presented in section 1 of the mathematical note in the SI. The analysis is performed in the limit *p, N → ∞* and *c* ≪ 1.

In our model, memories are defined as the patterns of external synaptic inputs that were present when the corresponding stimulus was shown for the first time to the network. These external synaptic inputs 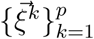, where *k* labels individual memories, are i.i.d. Gaussian random variables with zero mean and unit variance. These memories are imprinted in the connectivity matrix using the learning rule described in Equation 2. The firing rates *r*_*i*_(*t*) of neurons *i* = 1, *…, N* evolve according to the rate (Wilson-Cowan) equations (Wilson and Cowan, 1972), i.e. Eq. 1.

During the delay period, the external stimulus *I*_*i*_ is set to be zero. The steady states or fixed point attractors for the dynamics are given by the following set of nonlinear equations

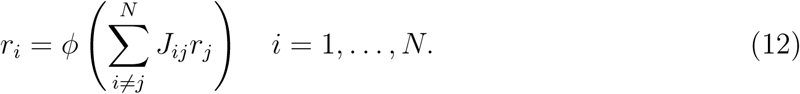

To describe the statistics of the firing rates in a fixed point described by Equation 12, we first need to compute the statistics of the incoming current to a given neuron, 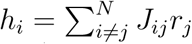, assuming that the network state is correlated with one of the stored patterns (without loss of generality, we choose here the first pattern 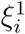), but uncorrelated with all other patterns. In the large *N* limit, the distribution of this current, conditioned on the value of 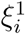, becomes a Gaussian. The mean *μ* conditioned on 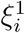 is given by

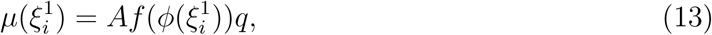

where *q* is the covariance between a non-linear transformation of the pattern 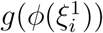 and the firing rates in the current network state *r*_*i*_,

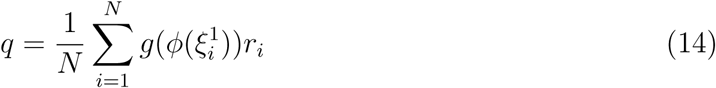

The ‘overlap’ *m* described in the main text is the corresponding correlation coefficient,i.e. normalized by the square root of the variances of 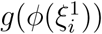 and *r*_*i*_.

The variance of input currents (due to the other stored patterns that act as a quenched source of noise on the retrieval of the pattern of interest) is given by

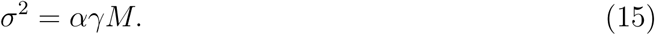

Where *γ* depends on the learning rule and statistics of the patterns as

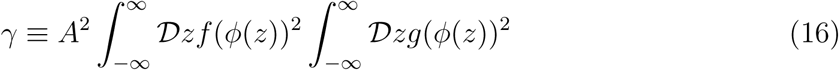

while *M* is the average squared firing rate

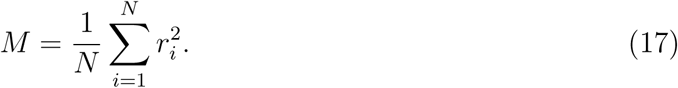

The next step is to compute self-consistent equations for the ‘order parameters’ *q* and *M*, that fully describe the macroscopic behavior of the network. Inserting Eq. (12) in Eq. (14),using the fact that *h*_*i*_ has a Gaussian distribution with mean *μ* and variance *σ*^2^, and replacing the sum over *i* by an integral over *ξ*_*i*_, we obtain

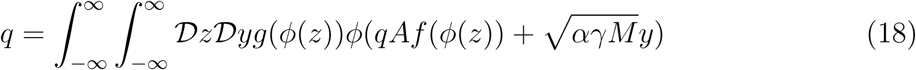

where the integral over *z* corresponds to an integral over the distribution of the patterns 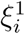, while the integral over *y* corresponds to an integral over the distribution of the ‘quenched noise’ due to other stored patterns.

Similarly, inserting Eq. (12) in Eq. (17) and using again the fact that *h*_*i*_ is Gaussian distirbuted, we find

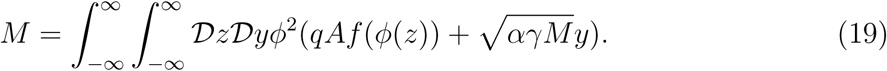

For a given value of *α*, and functions *ϕ*, *f* and *g*, Eqs. (18,19) are solved numerically by using a gradient free approach where the equations are iterated as a discrete map from an arbitrary initial condition (i.e. *q*_0_ *>* 0 and *M*_0_ *>* 0) until convergence. Note that Eq. (18) always have a solution *q* = 0, which correspond to a background state which is uncorrelated with all stored patterns. Solutions of these equations with *q >* 0 indicate the presence of retrieval states.

The distribution of firing rates can be obtained as

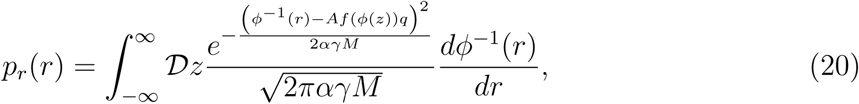

where the order parameters *q* and *M* are determined by the self-consistent equations (19) and (18). The overlap *m* is given by

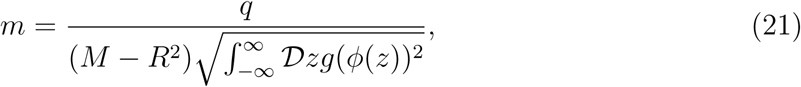

where *R* is the mean firing rate in the attractor state given by

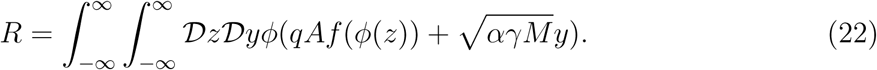

Similar analysis can be performed in the presence of an external input (presentation period), for both familiar and novel stimuli. Details are presented in section 2 of the mathematical note in SI. The calculations proceed along the lines of the calculations presented above, except that (1) During the presentation of a familiar stimulus, the external input currents are set to 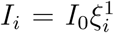; (2) During the presentation of a novel stimulus, the external inputs are set to 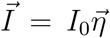 where 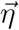 is an independent and identical distributed standard normal pattern of currents (i.e. 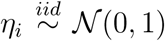 with *i* = 1, 2, *…, N*), uncorrelated with all learned patterns. For simulations in Figures 3 and 6 *I*_0_ was set equal to one during the presentation period.

### Simulations

For most simulations shown in this paper, the probability of connections was set to 0.5% (i.e. *c* = 0.005) and the number of neurons to *N* = 50000, which implies an average number of connections per neuron of *Nc* = 250. To investigate finite size effects, we also performed simulations with various values of *N* and *c* (see Fig. S5). The single neuron time constant was chosen as *τ* = 20*ms*, similar to time constants of single neurons (McCormick et al., 1985) and synapses (Destexhe et al., 1998), and with the decay time constant of cortical activity as measured *in vivo* (Reinhold et al., 2015). Open source built-in linear algebra methods in scipy and numpy Python packages suited for sparse matrices were used to generate the connectivity matrix. For simulating the networks dynamics, the Euler method was used with a time step size of 0.5ms. For a few parameter sets, we checked that results are unchanged when a smaller value of *dt* = 0.1ms is used. The code is available upon request.

## Acknowledgements

We thank Yonatan Aljadeff and Yali Amit for useful discussions and comments on previous versions of the manuscript. We also thank David Sheinberg for providing his data; Sukbin Lim for providing data analysis codes, and useful comments on the manuscript; and Maxwell Gillett for useful discussions on optimizing network simulations.

